# Chlorinated electron acceptor availability selects for specific *Dehalococcoides* populations in dechlorinating enrichment cultures and in groundwater

**DOI:** 10.1101/175182

**Authors:** A. Pérez-de-Mora, A. Lacourt, M.L. McMaster, X. Liang, S.M. Dworatzek, E.A. Edwards

## Abstract

Individual *Dehalococcoides mccartyi (Dhc)* strains differ primarily from one another by the number and identity of the reductive dehalogenase homologous catalytic subunit A (*rdhA*) genes contained within their respective genomes. While thousands of *rdhA* genes have been sequenced, the activity of the corresponding proteins have been identified in only a handful of cases. Most effort has focused on identifying the enzymes that dechlorinate substrates including trichloroethene (TCE), cis-dichloroethene (cDCE) and vinyl chloride (VC) relevant to groundwater remediation. The associated *rdhA* genes, namely *tceA, bvcA,* and *vcrA*, along with the *D. mccartyi* 16S rRNA gene are often used to track growth and dechlorinating activity in DNA extracted from field samples. In this study, we augmented the typical suite of three characterized *rdhA* genes to include an additional 12 uncharacterized *rdhA* sequences identified in the metagenome in the mixed *Dhc*-containing culture KB-1 to track population shifts within the culture and at two bioaugmented field sites. Quantitative PCR assays were developed for the 15 selected *D. mccartyi rdhA* genes and evaluated using 11 different sub-cultures of KB-1, each enriched on different chlorinated ethenes and ethanes. The proportion of *rdhA* gene copies relative to *Dhc* 16S gene copies indicated the presence of multiple distinct *Dhc* populations in each culture. The specific electron acceptor amended to each culture had a major influence on the distribution of *D. mccartyi* populations and their associated *rdhA* genes. We also surveyed the abundance of *rdhA* genes in samples obtained from two bioaugmented field sites. Growth of the dominant *D. mccartyi* population in the KB-1 inoculum was detected in the UK site samples. At both field sites, the measurement of relative *rdhA* abundances revaled significant *D. mccartyi* population shifts over time as dechlorination progressed from TCE through cDCE to VC and ethene, indicating that the selective pressure of the most abundant chlorinated electron acceptor that was observed in lab cultures was also occurring in the populations in the field. Understanding driving forces behind *D. mccartyi* population selection and activity is improving predictability of remediation performance at chlorinated solvent contaminated sites.

## Importance

Reductive dechlorination, often mediated by bacteria from the genus *Dehalococcoides*, is a very effective and now widely used method for bioremediation. These organisms couple growth to detoxification of chlorinated industrial solvents that contaminate groundwater. A better understanding of the breadth of populations within this genus and the conditions that lead to successful growth and complete dechlorination are critical to more widespread adoption of this cost-effective remediation strategy. This study identified environmental conditions that lead to the selection of different populations of *Dehalococcoides* that impact remediation performance.

## Introduction

There are thousands of public and private sites with chlorinated solvent related groundwater contamination problems (1). Chlorinated volatile organic compounds (cVOCs) such as tetrachloroethene (PCE) and trichloroethene (TCE) as well as their daughter products including the isomers of dichloroethene (DCE) and vinyl chloride (VC) are highly toxic compounds and TCE and VC are recognized human carcinogens by the National Toxicology Program (http://ntp.niehs.nih.gov/pubhealth/roc/roc13/). Clean-up of groundwater contaminated with these compounds takes time and is costly. Bioremediation and bioaugmentation have gained significant acceptance as viable approaches for treatment of chlorinated ethenes in the subsurface (2). The primary biotransformation mechanism for chlorinated ethenes in groundwater is reductive dechlorination under anaerobic conditions, which involves a stepwise replacement of Cl atoms with H atoms following the sequence: PCE, TCE, DCE (mainly cis-DCE), VC and finally non-toxic ethene (3-6).

Diverse anaerobic microorganisms (e.g., *Dehalococcoides*, *Desulfitobacterium, Dehalobacter*, *Sulfurospirillum, Desulfuromonas, Geobacter, Dehalogenimonas*) can couple reductive dechlorination of chlorinated ethenes with growth, a process called organohalide respiration (7-12). Nonetheless, dechlorination beyond DCE and VC has only been shown so far for members of the *Dehalococcoidales* (3, 5, 13-16). In practice, owing to subsurface heterogeneity, natural reductive dechlorination is incomplete in some locations, resulting in the accumulation of the daughter products cDCE and the carcinogen VC (17). This is generally attributed to poor mixing, lack of appropriate *D. mccartyi* organisms or electron donor, or inhibition of terminal dechlorination steps (2).

Bioaugmentation with mixed cultures containing *D. mccartyi* can overcome stalling or reduce the time required to achieve complete reductive dechlorination (2, 18-22). The abundance of *D. mccartyi* in groundwater is most often assessed via real-time qPCR of the 16S rRNA gene (23-25). While little dechlorination of cDCE and VC has been measured in the absence of *D. mccartyi*, the presence of detectable *D. mccartyi* does not always indicate that complete dechlorination to ethene will occur. The dechlorinating ability of *D. mccartyi* is strain-specific and is determined by the strain’s complement of reductive dehalogenase genes and their activity. Thus, *D. mccartyi* strains with identical 16S rRNA may differ in the chlorinated compounds that they can respire. Reductive dehalogenase enzymes (RDases) catalyze the carbon-halogen bond cleavage reaction, and offer an additional biomarker for tracking *D. mccartyi* strains. RDases are heterodimeric, membrane-bound enzymes, consisting of a catalytic active A unit of about 500 amino acids (aa) anchored outside of the cytoplasmic membrane by a small (100 aa) predicted integral membrane B subunit, encoded by *rdhA* and *rdhB* genes (26). Further elucidation of the respiratory electron transport chain of *D. mccartyi* suggests a multi-protein reductive dehalogenase complex consisting of the RDase, a molybdoenzyme and a hydrogen uptake hydrogenase (27). Due to their hydrophobic nature, oxygen sensitivity and complex association, only a few RDases have been biochemically characterized to date. Among these are the enzymes catalyzing the conversion of PCE to cDCE and TCE to VC and coded by the *pceA* and *tceA* genes, respectively, as well as the RDases catalyzing the conversion of cDCE to ethene coded by the *bvcA* and *vcrA* genes (28-32). Quantitative PCR methods that target these specific genes have been developed and are being increasingly used as prognostic and diagnostic tools in the field to overcome the limitations of the 16S rRNA gene (33-35).

The genomes of more than 10 *D. mccartyi* isolates have now been sequenced. These genomes are highly streamlined (∼1.4 Mb) and striking in their similarity, differing primarily in two regions termed High Plasticity Regions (HPR) on either side of the ORI. Each genome harbors many distinct full-length reductive dehalogenase homologous genes (*rdh*AB) per genome (*e.g.,* 17 in strain 195, 32 in strain CBDB1 and 36 in strain VS) (36-38). Hundreds if not thousands more putative *rdhAB* genes have been identified from metagenome sequencing efforts. Owing to the lack of functional characterization for most of this protein family, a sequence identity-based classification of orthologues into groups based on >90% a.a. identity was developed (39). This sequence-based classification was adopted prior to having a crystal structure to identify active site and other key residues. Fortunately, the two crystal structures recently solved (40, 41) support the original classification, and the database of sequences and new ortholog groups continues to expand (39, 66).

In this study, we attempted to quantify a larger suite of *rdhA* genes to help distinguish different *D. mccartyi* populations from each other in mixed cultures and groundwater samples where multiple closely related populations may coexist. We first compared the *rdh*A complement of the mixed culture KB-1 with those of eleven isolated *D. mccartyi* strains to identify *rdhA* sequences (and their corresponding Ortholog Groups) that are less commonly shared between strains. Methods for quantitative real-time PCR were developed for the selected *rdh*A genes and these assays were first tested in subcultures of KB-1 enriched on different chlorinated terminal electron acceptors. Finally, we used the selected suite of *rdh*A genes to detect the presence and dynamics of different *D. mccartyi* populations in groundwater from two bioaugmented sites.

## Materials and Methods

### Cultures and growth conditions

The KB-1 consortium is a functionally-stable enrichment culture that originated from microcosms prepared from aquifer material from a TCE-contaminated site in southern Ontario (4). KB-1 is routinely maintained in batch mode with TCE as electron acceptor, and dechlorinates PCE through TCE, cis-DCE and VC to ethene. A transfer from the original KB-1 culture has been grown and used commercially for more than a decade for bioaugmentation at cVOC-contaminated sites (siremlab.com). The main organisms in the KB-1 culture have been investigated over the years via clone libraries, qPCR quantification and metagenome sequencing (42-47). Two dechlorinating genera have been identified in the culture, namely *Dehalococcoides* and *Geobacter,* that are supported by many other fermenters, acetogens and methanogens (44, 48). Many years ago, the original TCE-fed KB-1 enrichment culture was used to inoculate various sub-cultures maintained on different terminal electron acceptors (TCE, cDCE, VC and 1,2-dichloroethane [1,2-DCA]). These subcultures have been maintained with methanol (M), hydrogen (H_2_), or a mixture of methanol and ethanol (ME) as electron donor. Table S1 summarizes the main features and growth conditions of all the different cultures studied with the name format indicating electron acceptor amended/donor used and year created. The commercial KB-1^®^ culture is referred to as TCE/ME_2001_SiREM in Table S1. All cultures were grown anaerobically in a defined minimum mineral medium (4). Cultures maintained at the University of Toronto are grown either in 0.25L bottles sealed with screw caps with mininert valves or in 1 or 2 L glass bottles sealed with black butyl stoppers. Typically, bottles contained 10% by volume of headspace flushed with a N_2_/CO_2_ 80%/20% as needed. These cultures are kept in the dark in an anaerobic glovebox at room temperature (22-25 °C). At SiREM, KB-1^®^ is grown in 100 L stainless steel vessels at 22-25 °C. At the University of Toronto, dechlorinating cultures were typically re-amended every 2-3 weeks. Cultures maintained at SiREM are re-amended more frequently, typically every 3 to 4 days, as substrate is depleted.

### KB-1 Metagenome data and *rdh* sequences

The parent KB-1 culture has been maintained at the University of Toronto since 1998 with TCE as the electron acceptor and methanol as the electron donor. This culture is referred to as “TCE/M_1998_Parent” in Table S1 (Supporting Information) and all other KB-1 enrichments originated from this parent culture. The partially assembled metagenome sequence of TCE/M_1998_Parent is publically available at the Joint Genome Institute (JGI) (http://genome.jgi-psf.org/aqukb/aqukb.download.ftp.html). Details on the extraction of genomic DNA for sequencing and the assembly of the KB-1 metagenome and of a draft chimeric genome *of D. mccartyi* strains are provided elsewhere (44, 49). A total of 31 distinct *rdh*A gene sequences have been identified in the KB-1 cultures from multiple investigations (47, 50). Thirty sequences are associated with *D. mccartyi*, whereas one sequence is from the *Geobacter* population present in KB-1. Table S2 (supporting information) compiles all previously identified KB-1 *rdhAB* sequences, which include the original 14 sequences found by Waller *et al*. in 2005 (51) as well as those identified more recently from metagenome sequencing (52) where an additional 13 *rdhAB* full sequences and two *rdhA* partial sequences were annotated (39, 50). For ease of reference and consistency, these additional *rdhA* sequences have been renamed and are now deposited to Genbank under the accession numbers KP085015-KP085029 to replace the previous JGI gene locus tags (Table S2).

### Phylogenetic analysis of *rdh*A sequences

A total of 249 *D. mccartyi rdh*A sequences from sequenced and characterized isolated strains were retrieved from NCBI for phylogenic comparisons. These sequences include available sequences from eleven isolated and sequenced *D. mccartyi* strains, an incomplete set of 3 *rdhA* sequences from strain MB (available at the time of retrieval), and the *rdh*A sequences found in KB-1. The number of *rdhA* sequences contributed by each strain was as follows: 195 (17), Bav1 (10), BTF08 (20), CBDB1 (32), DCMB5 (23), FL2 (14), GT (20), GY50 (26), JNA (19), MB (3), VS (37), and KB-1 (28 [*Dhc*]+1 [*Geobacter*]). Two of the KB-1 *D. mccartyi rdhA* sequences and sequence VS_1308 are only partial sequences (less than 850 nucleotides) and were therefore not included in phylogenetic comparisons. The details of alignments and tree construction are provided in Supporting Information (Text S1). Files containing all amino acid and nucleotide sequences as well as matching of tree nomenclature with protein names can be found in a folder labelled rdhA_Dhc_seq150420 at the following address: https://docs.google.com/folder/d/0BwCzK8wzlz8ON1o2Z3FTbHFPYXc/edit.

### Groundwater sampling and site description

Groundwater was collected from two TCE-contaminated sites prior to and after bioaugmentation with KB-1^®^. The first site was located in Southern Ontario (ISSO site) and was characterized by contaminants in fractured bedrock. Figure S1a (Supporting Information) shows details of relevant sampling dates and events related to this site. Samples corresponding to three different phases of the remediation were investigated: predonor or pretreatment phase, ii) biostimulation phase, consisting of the daily addition of ethanol as an electron donor and iii) bioaugmentation phase, following the inoculation of KB-1^®^ (Figure S1a). At the ISSO site, a groundwater recirculation system consisting of two injection and three extraction wells was installed to improve the distribution of electron donor (ethanol) and microorganisms (Figure S1b). Groundwater from the extraction wells was combined into a central manifold (composite), filtered, treated with chlorine dioxide (ClO_2_) to control biofouling, and amended with electron donor (ethanol) prior to distribution into the individual recharge wells. During both the biostimulation and bioaugmentation phases, ethanol was added on a daily basis. Bioaugmentation with KB-1^®^ consisted of one single addition of approximately 100L of culture. Groundwater samples for molecular analysis were obtained from the composite pipeline, where groundwater from the three extraction wells was combined (Figure S1b). Additional information on this site can be found in a previous publication that surveyed community dynamics at the site (22).

The second site is located in the U.K. and consisted of a pilot test cell (30x7x4 m) for treatment of a DNAPL source area (*∼* 1000 kg of DNAPL within the cell). The cell was conceived as an “in situ laboratory” for investigating source area bioremediation (SaBRE project - http://www.claire.co.uk/index.php?option=com_content&view=article&id=53&Itemid=47). The relevant sampling dates and events related to this site and a sketch with sampling locations are shown in Figures S2a and S2b (Supporting Information). Samples for investigation were collected: i) prior to any remediation action; and ii) after donor addition of one single dose of donor SRS^™^, a commercially available emulsified vegetable oil (Terra Systems, Inc.) and bioaugmentation with KB-1^®^ (Figure S2a). Groundwater was collected from fully screened sampling wells (SW) at four locations within the test cell: i) at the influent (INF); within the source zone (SW70); iii) within the plume (SW75); and iv) at the effluent (EFF), also within the plume. The test cell was operated initially for a 90-day baseline period to establish steady-state pre-treatment conditions. Groundwater was extracted at an average of 1.4 liters per minute, corresponding to an average residence time within the cell of 45 days. A total of 2,400 Kg of SRS^™^ at a 5% concentration was used as the electron donor and injected along the test cell. Two weeks after donor injection approximately 65 L of KB-1^®^ was added using the same injection ports that were used to add electron donor. Within the test cell, both TCE and cDCE were the main cVOCs. A dissolved phase plume emanating from the source and extending more than 400 m away was further characterized by the presence of VC and ethene. The starting point (Day 0) was defined as the day following the end of the emulsified oil injection, which took approximately one week to complete. Further details on the SaBRE site can be found in technical bulletins freely available on the SaBRE-CL:Aire website (see above).

### Nucleic Acid Extraction

Samples were collected from eleven different enrichment cultures at one or two times each during 2011 (Table S1). Archived samples from the TCE/ME_2001_SiREM culture spanning 5 years were obtained from SiREM. DNA was also extracted from groundwater samples at from the two field sites described above. For extraction and isolation of genomic DNA (gDNA), samples from cultures (10-50 mL) or groundwater (200-1000 mL) were filtered through Sterivex^™^ (Millipore, MA) (0.22 μm pore size) filters using a centrifugal pump and dual-trap system. Filters were subsequently stored at −80 °C until further processing. For gDNA extraction, the casing of the filter was opened and the filter cut in about 30 pieces of similar size. The latter were introduced into a 2 mL nucleic acid extraction tube containing buffers and beads (Mo Bio Laboratories UltraClean® Soil DNA Isolation Kit, CA) and DNA extraction was completed following the manufacturer’s instructions. Elution of nucleic acids from the silica membrane was performed using 50 *μ*L of UltraPure^™^ DNase/RNase-Free distilled water (Invitrogen, CA) for samples from cultures as well as from the ISSO site in Canada, whereas 920 *μ*L were used for elution of samples from the SaBRE site in the UK.

### Quantitative PCR (qPCR) Amplification of Extracted DNA

Quantification of the 16S *rRNA* gene of *Dehalococcoides* and *Geobacter*-KB1 as well as a total of 15 *D. mccartyi rdhA* genes plus one *Geobacter rdh*A gene from the KB-1 culture was achieved via real-time qPCR on an Opticon 2 thermocycler (MJ Research). Table S3 (Supporting Information) provides details on the primer pairs used, their annealing temperatures and the length of the amplicon generated. With the exception of five genes (KB1-6/*bvcA*, KB1-14/*vcrA,* KB1-27/*tceA*, KB1-1 and KB1-5), new primers were designed for all other *rdhA* genes in this study. The specificity of the probes was tested *in silico* against the public NCBI NR nucleotide database and tested *in vitro* against DNA extracted from mixed dechlorinating and non-dechlorinating cultures. Table S4 (Supporting Information) shows the number of mismatches for each primer pair with sequences belonging to the ortholog group of the targeted *rdhA* gene. The PCR reaction (20 μL) consisted of: 10 μL of SsoFast^™^ EvaGreen® Supermix (Biorad, CA), 0.5 μL of each primer (10 mM), 7 μL of UltraPure^™^ DNase/RNase-Free distilled water (Invitrogen, CA) and 2 μL of template. The amplification program included an initial denaturation step at 98 °C for 2 min followed by 40 cycles of 5 s at 98 °C and 10 s at the corresponding annealing temperature. A final melting curve from 70 to 95 °C degrees at increments of 0.5 °C per second was performed. Since two concentrations were tested per template (undiluted and 10-fold diluted) for assessment of potential matrix-associated inhibitory effects, reactions for each dilution were performed in duplicate. The undiluted sample generally contained between 10-20 ng of DNA per μL as measured using a Nanodrop spectrophotometer (Thermo Scientific, DE). Generally, there was good agreement between the two measurements. Ten-fold serial dilutions of plasmid DNA containing one copy of the 16S *rRNA* gene or the *rdhA* gene (∼1000-1500 bases) were used as calibrators. Plasmids for calibration were prepared by PCR amplifying the desired *rdh*A from KB-1, inserting the gene into the pCR 2.1 PCR vector and subsequently transforming the construct into TOP10 *E. coli* competent cells (Invitrogen, CA) as per the TOPO TA cloning^®^ kit (Invitrogen, CA). Plasmid DNA was purified using the GenElute^™^ Plasmid Miniprep Kit following the manufacturer’s instructions (Sigma-Aldrich, MO). DNA was eluted using 50 μL of Ultra Pure DNase/RNase-free distilled water (Invitrogen, CA). Standard curves exhibited linear behavior (R^2^>0.990) when plotted on a logarithmic scale over seven orders of magnitude. Table S5 (Supporting Information) provides the details on the standard curves for each primer pair, including slopes (efficiencies), Y-intercepts and calibration model fit. The specificity of the amplicons was checked by melt curve analysis as well as by agarose gel electrophoresis for selected samples. Non-template controls were included in each run. Absolute gene copy concentrations for all cultures and field samples are provided as supporting information (Tables S6-S8; Supporting Information). The method detection limits (MDLs) were expressed in terms of gene copies per L of groundwater or per mL of culture and varied depending on the volume of sample filtered and the volume employed for elution of DNA from the purification column. The MDLs for each set of samples are provided in Table S9 (Supporting Information).

### Cluster analysis of *rdh*A sequences and calculation of *rdh*A/16S rDNA *Dhc* ratios

Phylogenetic analysis of *rdh*A nucleotide and amino acid sequences, including bootstrapping, was performed with the ClustalX free software (version 2.0.12; University College Dublin, [http://www.clustal.org/clustal2/]). Protein sequences were clustered into orthologue groups as previously defined (90% pairwise identity in amino acid alignments; per (39)). The similarity of the *Dehalococcoides* populations in the various KB-1 subcultures was investigated on the basis of their *rdhA* fingerprints (*rdhA*/16S rDNA *Dhc* ratios) by means of hierarchal cluster ordination analysis. The latter was performed with the free software Hierarchal Clustering Explorer (HCE) (version 3.5; Human-Computer Interaction Lab, Univ. of Maryland, [www.cs.umd.edu/hcil/hce/]). Details on the elaboration of the phylogenetic trees and clustering analysis are provided in the Supplementary information (Text S1 and Text S2).

## RESULTS AND DISCUSSION

### Reductive dehalogenase (*rdhA*) genes in the KB-1 consortium

We first thought to distinguish *D. mccartyi* populations in our cultures using unique non-coding regions from metagenomic sequences, but the core genome is basically identical for all *D. mccartyi* populations in KB-1 we could not identify suitable regions for primer design. Thus, we turned to using a suite of *rdhA* genes for this task. To better understand the similarity between *rdhA* sequences, we constructed phylogenetic trees using 249 amino acid (Figure 1) and nucleotide (Figure S3; Supporting Information) sequences that included all full *rdhA* sequences found in the KB-1 metagenome as well as those identified in eleven isolated *D. mccartyi* strains. Amino acid sequences belonging to the same ortholog group (OG), defined as having > 90% PID (39), are highlighted in different colors on the phylogenetic trees. These highlighted branches clearly support the classification and reveal how some sequences are present in all strains, while others only in a few. When the same sequences are compared at the nucleotide level (Figure S3) identity within an ortholog group can be substantially lower than 90%. The terminology reductive dehalogenase *homologous* genes (*rdhA*) was adopted years ago, and perhaps implies more knowledge of common ancestry than is in fact known, but it is not unreasonable to suspect that these genes arose from either speciation (orthologous) or duplication (paralogous) events. Herein we use the term “ortholog” to refer to *rdhA* genes that group together according to the classification system proposed by Hug et al. (39) with >90% pairwise amino acid identity. The term “homologous” is used to refer to all *rdhA* genes regardless of which ortholog group they belong to.

**Figure 1.**
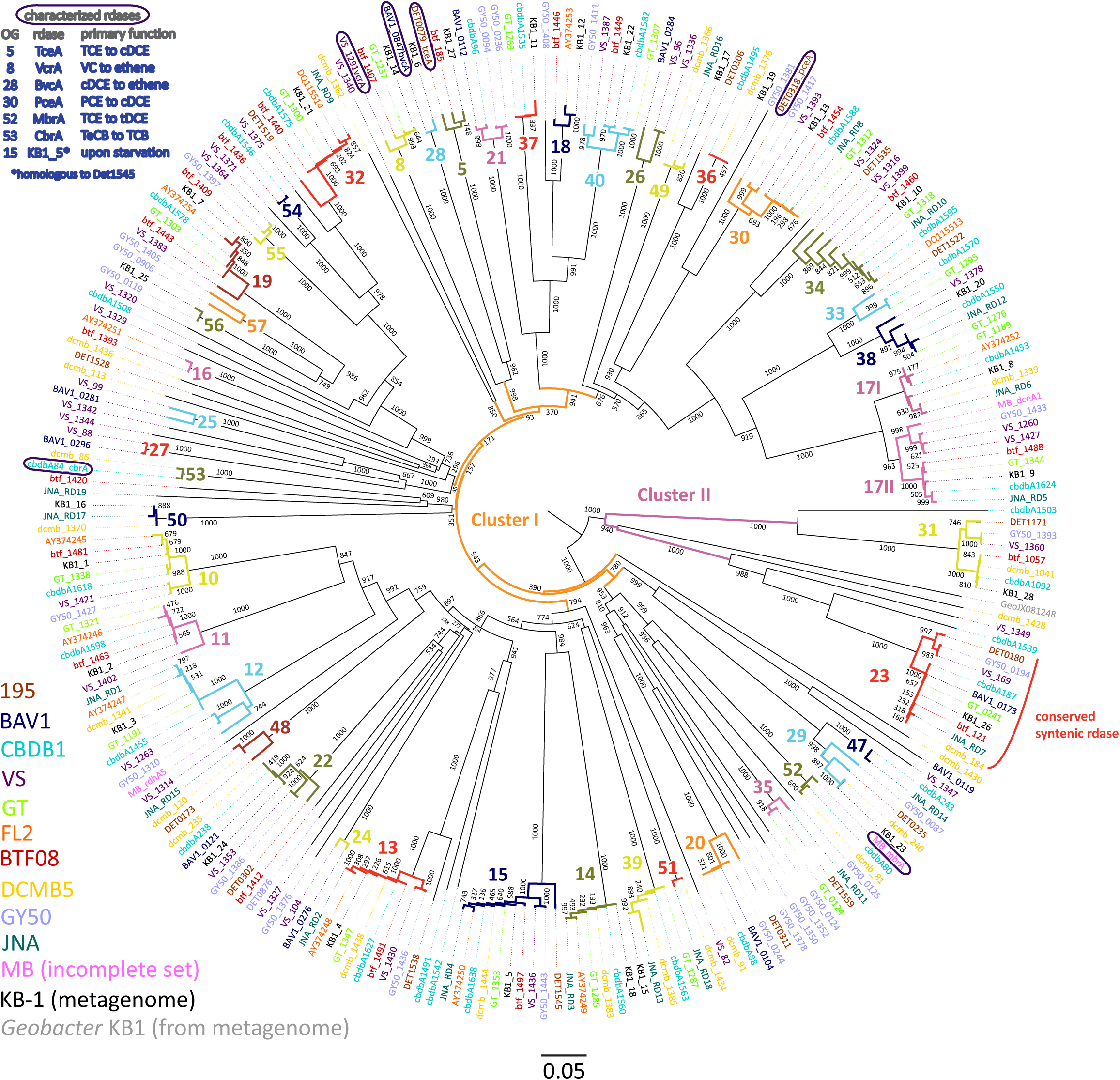
Phylogenetic tree of *D. mccartyi* reductive dehalogenases based on amino acid sequences. Tree includes sequences predicted from 248 *rdhA* genes found in eleven isolated *D. mccartyi* strains as well as those found in the mixed dechlorinating culture KB-1, including the *rdhA* gene sequence found in *Geobacter* KB-1 as an outgroup. The scale is the number of susbtitutions per amino acid. Bootstraps are shown at tree nodes. Sequences names are colored corresponding to their source as shown in the legend at bottom left. The numbers and different branch colors highlight ortholog groups (OGs) where sequences have > 90% PID. Sequences corresponding to characterized RDases are circled and their substrates shown in the top right legend (chlorinated substrate abbreviations as in text, plus TeCB: tetrachlorobenzene; TCB: trichlorobenzene).

The phylogenetic analysis of 249 *D. mccartyi rdh*A sequences generated a total of 43 ortholog groups (OG) including 6 new previously un-described groups (OG 52 to 57). Of the 249 sequences analyzed in 2016, 37 sequences remained ungrouped, meaning without a single ortholog (Figure 1). At the amino acid level, none of the KB-1 *D. mccartyi rdhA* sequences was unique because each *rdhA* gene in KB-1 has more than 90% amino acid sequence identity with at least one other *rdhA* sequence in another *D. mccartyi* strain. Indeed, most *rdhA* sequences in KB-1 were found to be associated with ortholog groups comprising multiple *rdhA* sequences (Figure 1). The existence of numerous shared *rdhA* genes among *D. mccartyi* is consistent with their co-localization with insertion sequences and other signatures for horizontal gene transfer, and within genomic islands in high plasticity regions (37).

### Selection of suite of distinguishing KB-1 *rdh*A genes

We selected a total of 15 characterized and uncharacterized rdhA sequences to track using qPCR. Ten uncharacterized *rdhA* sequences, KB1-25 (ortholog group (OG) #56), KB1-11 (OG37), KB1-12 (OG18), KB1-16 (OG50), KB1-17 (OG 49), KB1-19 (OG36) and KB1-23 (OG29), were selected arbitrarily as those with the fewest orthologs in other strains based on our tree (Figures 1 and S3). KB1-15 (OG39) and KB1-18 (OG14) were selected because they were found on the same contig in the KB-1 metagenome and both share homology to genes in strains GT, CBDB1, DCMB5 and JNA, therefore they may be mobilized together (Figure 1). We included KB1-4 (OG13) because it appears to have orthologs in all other strains and could be useful for normalization. To this set of 10 uncharacterized genes, we added five more genes of interest: KB1-1 (OG10), KB1-5 (OG15), KB1_14/*vcrA* (OG8), KB1_6/*bvcA* (OG28) and KB1_27/*tceA* (OG5). KB1-1 and KB1-5 were selected because their corresponding proteins have previously been detected in KB-1 (54, 55). Furthermore, KB1-5 (OG 15) is orthologous to DET1545 from Strain 195, which is expressed upon starvation (55, 56). KB1_14, KB1_6 and KB_27 correspond to the functionally-characterized vinyl chloride and trichloroethene reductases VcrA, BvcA and TceA. We designed qPCR primers to these 10 uncharacterized genes, and used previously designed primers for remaining genes (Table S4). Next, we monitored the abundance of this suite of rdhA genes in DNA samples from 11 different KB-1 enrichment sub-cultures maintained over years on different chlorinated electron acceptors (Table S1).

### Quantification of *rdh*A genes in KB-1 enrichments with different chlorinated electron acceptors

To reflect shifts in *D. mccartyi* populations, qPCR results are presented as *rdhA* gene copies divided by 16S rRNA gene copies to provide an approximation of the relative proportion of each *rdhA* gene per *D. mccartyi* genome. *D. mccartyi* are known to harbor only one copy of the 16S *rRNA* gene per genome. *RdhA*/16S rRNA ratios were visualized using heatmaps with values ranging from above 0.6 (where more than 60% of all *D. mccartyi* genomes in the culture contains that gene) to less than 0.001 (where the gene is present in fewer than 0.1% of the *D. mccartyi* populations) (Figure 2). We color-coded these ratios (x) into one of five abundance categories (x≥0.6 (dark red); 0.6 >x≥0.1 (red); 0.1>x≥0.01 (orange); 0.01>x≥0.001 (yellow) and x<0.001 (pale yellow)). Although the primers were designed to target specific KB-1 *rdhA* sequences, most sequences within the same ortholog group would likely also be amplified. Table S4 (Supporting Information) compiles all mismatches found between primers used and other sequences in the corresponding OG.

**Figure 2.**
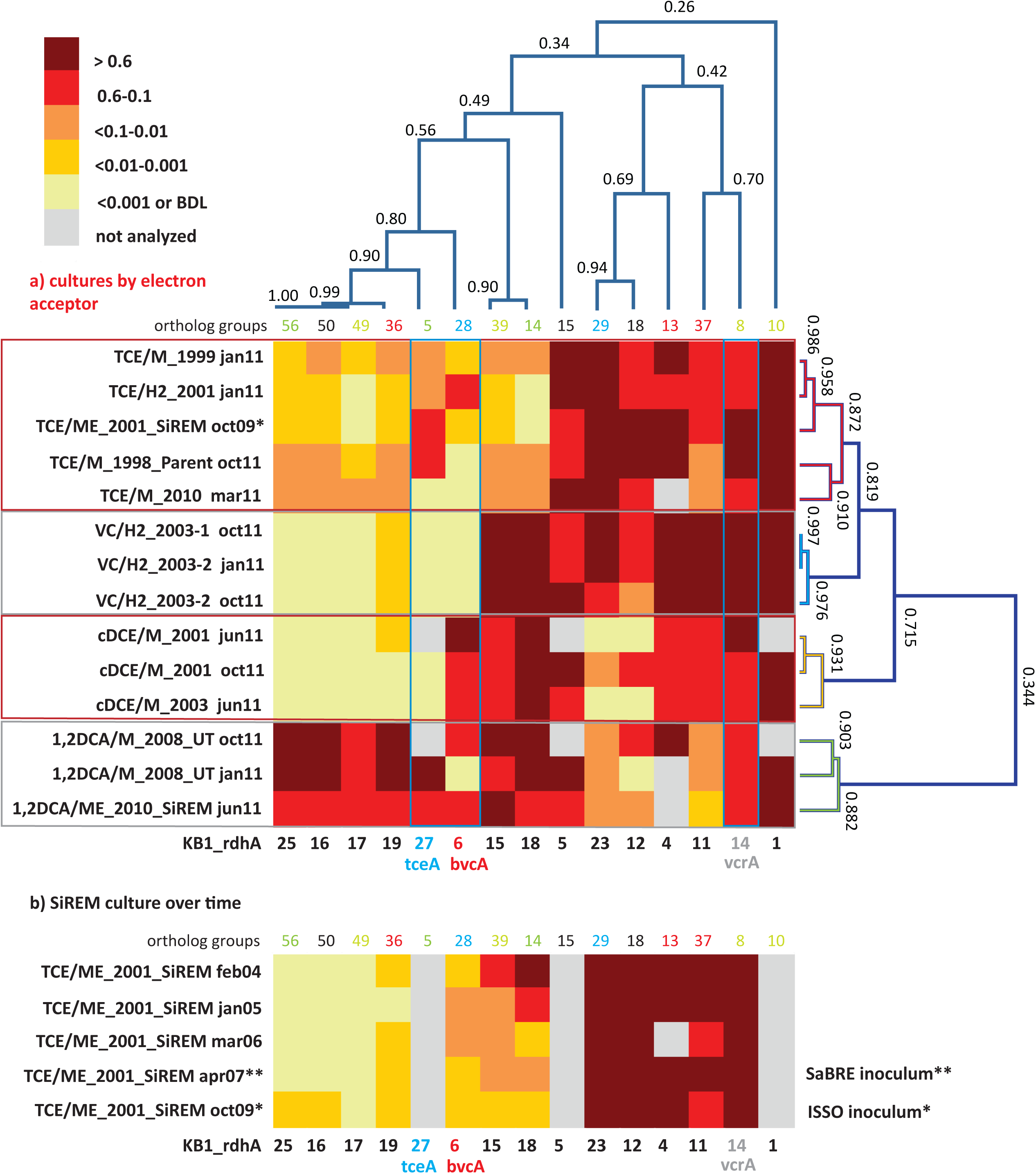
Cluster analysis and heatmaps showing *D. mccartyi rdhA*/16S rDNA gene copy ratios measured in KB-1 enrichment cultures. Panel a) shows how data clustered by the chlorinated electron acceptor amended to individual enrichment cultures as well as by *rdhA* sequence. The name format indicates electron acceptor amended/donor used_year created, followed by date sampled (e.g., TCE/M_1999 jan11 is a TCE and methanol enrichment culture first established in 1999 and sampled in January of 2011). The numbers on the cluster branches of both axes indicate the percentage similarity between samples based on the Pearson’s correlation coefficient. Panel b) is a heatmap of *rdhA*/16S gene copy ratios in the SiREM KB-1 culture over 5 years. Cultures indicated with * or ** are those used for bioaugmenting field sites in 2007 (SaBRE) and 2009 (ISSO).

*D. mccartyi rdhA*/16S rDNA gene copy ratios measured in KB-1 enrichment cultures (Figure 2) revealed more variability, and thus a greater number of distinct *D. mccartyi* populations than anticipated, considering the number of years of enrichment and the fact that they all originated from a common parent culture. Clustering analysis revealed rdhA genes common to all cultures and *rdhA* gene specific to a particular chlorinated electron acceptor. For instance, all subcultures were found to contain KB1-1 (OG10) (ratios all greater than 0.6), and to have consistently abundant *rdhA* sequences corresponding to KB1-4 (OG13) and KB1-5 (OG15) and KB1-14 (*vcrA*; OG8). The most striking finding (Figure 2a) is the clustering of specific patterns with terminal chlorinated electron acceptor. The VC enrichments show the lowest diversity of *rdhA* sequences and the most concordance between enrichments. These enrichments contain high abundances of KB1-1, KB1-14/*vcrA*, KB1-11, KB1-4, KB1-15 and KB1-18 with more variable but still abundant presence of KB1-5, KB1-23 and KB1-12. Other *rdhA* sequences are near or below detection. The cDCE enrichments form their own cluster, distinct from the VC enrichments because they contain significant abundance of the KB1-6/*bvcA* gene, and much lower proportions of KB1-23 and KB1-12. These cDCE enrichments were the only ones consistently enriched in *bvcA*. The 1,2-DCA enrichments also formed a distinct cluster characterized by high ratios for KB1-16, KB1-17, KB1-19, KB1-25 as well as KB1-27/*tceA*. Finally, the TCE cultures reflect a blend of all the other enrichments, showing more variability, although favoring a pattern most like the VC enrichments. The clustering analysis also revealed co-variation among some of the *rdhA* genes regardless of the enrichment, suggesting co-location of these genes on the same genome as is the case of KB1-15 and −18, and possibly also KB1-16, −17, −19 and −25 (Figure 2a). The *Geobacter rdhA* gene was detected at similar abundance to the *Geobacter* 16S rRNA gene only in the TCE-amended cultures (at approx. 15-25% of *Dhc* abundance); in all other enrichments *Geobacter* 16S rDNA and *rdhA* genes were below the detection limits (Tables S6 and S7), as expected since *Geobacter* only dechlorinates PCE or TCE as far as cDCE. In control DNA samples from cultures without *D. mccartyi,* copies of *rdhA* genes were all below the MDL (data not shown).

The stability of the *rdhA* fingerprints over time was also assessed. For the VC, 1,2-DCA and cDCE enrichments, two samples from the same enrichment culture bottle were analyzed at 4 or 9 month’s intervals (Jan and Oct or Jun and Oct, 2011; Figure 2a). There was good agreement between the two timepoints, although small differences likely reflect changes in relative abundance of *D. mccartyi* populations in batch incubation conditions. For the TCE/ME_2001_SiREM culture, five DNA samples over a time span of 5 years (2004, 2005, 2006, 2007 and 2009) were available, and were investigated using a subset of *rdhA* genes (Figure 2b). The *rdhA* fingeprint was relatively stable over time except for three *rdhA* genes, namely KB1-18, KB1-15 and KB1-6/*bvcA*. Ratios for KB1-18 and KB1-15 gradually shifted from greater than 0.6 in 2004 to less than 0.01 and even less than 0.001 by 2009. KB1-6/*bvcA* ratios fluctuated over this time period, increasing above 10% in 2005-06 and back down to less than 1% in 2004, 2007 and 2009. As noted already, KB1-18 and KB1-15 appear to co-vary, supporting their co-localization in the same genome. These two genes are abundant in VC, cDCE and 1,2-DCA enrichment cultures but not in the TCE enrichments and diminished over time in the TCE/ME_2001_SiREM. These data reveal that the dominant VC-dechlorinating, *vcrA-* containing *D. mccartyi* populations are not the same in the TCE and VC enrichment cultures. It appears that the dominant *D. mccartyi* population more recently present in the VC/H_2_ sub-cultures is more similar to the one that was originally present in the parent culture, and that this population has been gradually surplanted in TCE/ME_2001_SiREM culture. This data provides a timeframe (∼5 years) for a major shift to happen in the *rdhA* fingerprint of a mixed culture maintained consistently on the same electron acceptor. This is different to the wholesale changes observed when the electron acceptor is changed, where a shift in the dominant *D. mccartyi* population may happen much faster, as demonstrated in a recent studies by Mayer-Blackwell *et al.* (57, 58) where the dominant *D. mccartyi* population in a bioreactor shifted over a period of 50-100 days when the electron acceptor was switched for 1,2-DCA from TCE. In the field, where conditions are more variable both in time and space than in the laboratory, changes in *D. mccartyi* populations may occur even faster.

The clustered data for 15 *rdhA* genes across 14 DNA samples from 11 different cultures yielded four major groups corresponding to each of the four chlorinated electron acceptors (Figure 2a). To create a more visual representation of the distribution of these rdhA genes in the two High Plasticity Regions (HPRs) of *D. mccartyi* genomes, we mapped corresponding OG groups in the order and HPR they typically appear in published genomes (Figure 3). This map provides evidence not just of multiple strains but also of likely gene deletions. As indicated previously, the considerable variability in the *rdhA* to 16S ratios within each culture and electron acceptor group was at first very surprising, because the cultures all derive from the same parent and have been enriched for so long on the same substrate. We expected to see only one clearly dominant population, especially in the VC to ethene and 1,2-DCA to ethene cultures that involve only a single dechlorination step. What the data show instead is a more complex pattern in each group, suggesting the presence of more than one *D. mccartyi* population even in the single-dechlorination step, highly enriched VC/H_2_ cultures. We repeated many of the DNA extractions and qPCR reactions to verify results, with no appreciable change in results. Reanalyzing samples from the enrichment cultures on more recent DNA (data not shown; manuscript in prep) also has not changed the results. Early experiments by Duhamel *et al.* (4, 5) and Waller *et al.* (51) had identified at least two distinct *D. mccartyi* populations (KB1-PCE and KB1-VC), the former containing KB1-6 (*bvcA*) and the latter not; these two strains could actually also be distinguished by a single difference in their 16S rRNA sequences (5). Subsequently, Hug (47) was unable to close the assembly of a *D. mccartyi* genome from the KB-1 TCE/M_1998 parent culture metagenome, particularly in the high plasticity regions rich in *rdhA* genes, because of the presence of multiple, highly similar *D. mccartyi* populations. Considering the cluster diagram by enrichment culture (Figure 2a; right side), we can perhaps infer at least four distinct populations based on *rdhA* distribution in the TCE enrichments, at least two in the VC enrichments (one with and one without *rdhA* KB1-12 (OG18), at least two in the cDCE enrichments and three in the 1,2-DCA enrichments (Figure 3). In an attempt to verify these results, we sequenced metagenomes from representative VC, cDCE and 1,2-DCA enrichments. Sequencing has confirmed the presence of multiple distinct *D. mccartyi* genomes in all enrichment cultures, even in the VC enrichments, upholding these results (data not shown; manuscript in preparation). The co-existence of multiple *D. mccartyi* populations at different cell densities within these enrichments likely arises from subtle substrate preferences of the expressed reductive dehalogenases and competition for available nutrients and vitamins. Functional characterization of some reductive dehalogenases reveals substrate overlap yet specific substrate preferences as observed for BvcA and VcrA (55, 61). Moreover, availability and type of corrinoid can alter rates of dechlorination for certain enzymes (62). Low abundance populations may persist in these cultures because they are maintained in batch mode, with infrequent medium changes and thus long residence times from 30 to 100 days.

**Figure 3:**
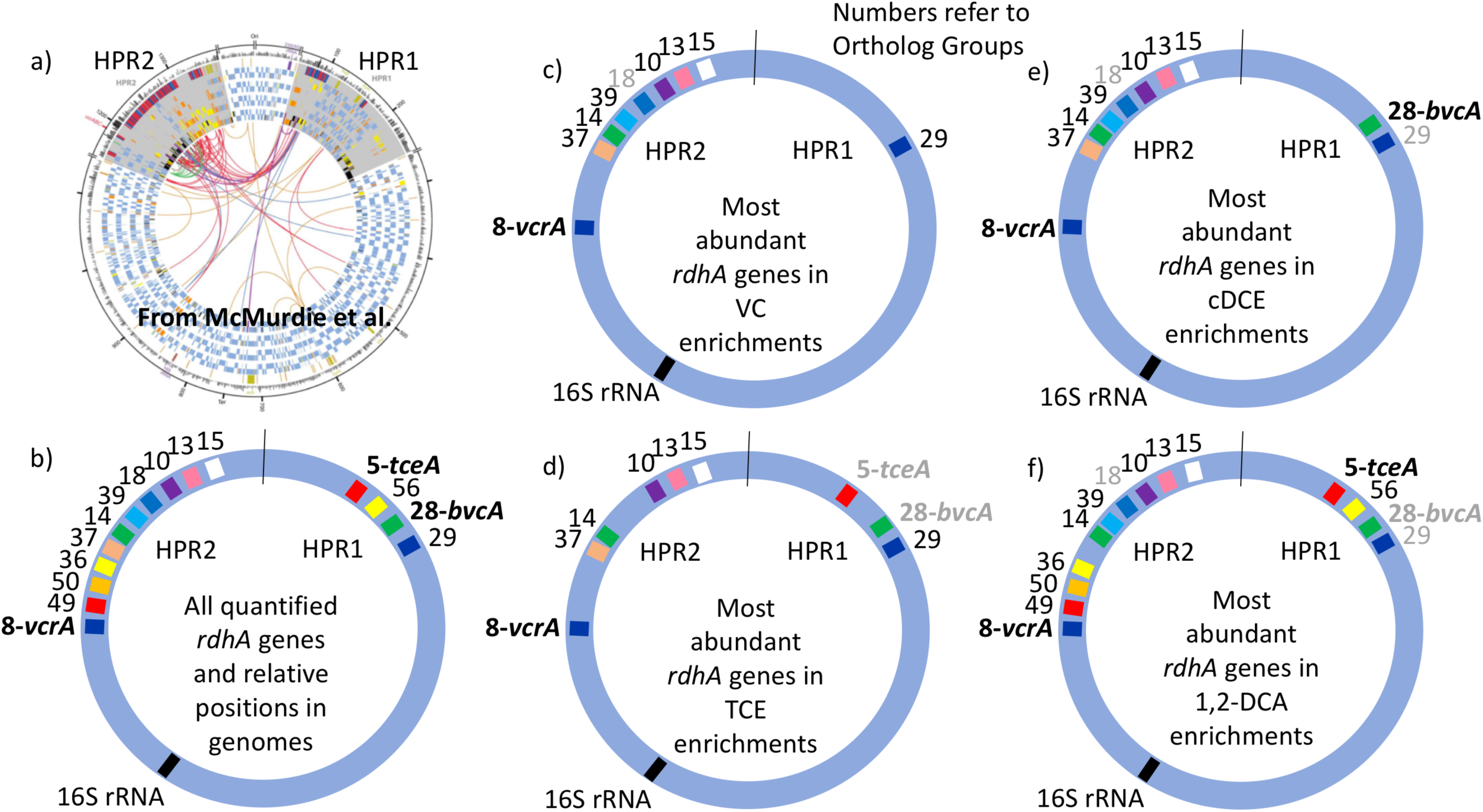
Distribution of most abundant *D. Mccartyi* reductive dehalogenase homologous (*rdhA*) genes quantified in KB-1 enrichment cultures. RdhA sequences are represented by their corresponding ortholog group (OG) numbers and relative positions in High Plasticity Regions (HPR) 1 and 2, as defined by McMurdie et al, 2009. a) Circular map of four *Dehalococcoides* genomes reproduced from McMurdie et al., 2009, illustrating the two high plasticity regions where most *rdhA* genes are found; origin is at top. b) Relative position of all of the rdhA sequences quantified by qPCR in this study. c) through f) most abundant *rdhA* genes detected in the respective enrichment culture samples, with evidence for considerable strain variation within and between enrichment cultures. OG numbers that are grey correspond to genes that are abundant in only some of the samples from the respective enrichments. See Table S3 or Figure 1 for correspondence between OG and KB-1 *rdhA* numbers.

### Quantification of *rdhA* genes in site groundwater

Because the KB-1 bioaugmentation culture harbors many dominant and minor populations of *D. mccartyi* in different abundances, monitoring the fate of these populations once added to a site became even more challenging than anticipated, as both the low and high abundance populations might grow and change in relative proportions, depending on the conditions and available electron acceptors at the site, just as they did in the enrichment cultures. A suite of 12 *rdh* genes was used to monitor *D. mccartyi* populations before and after bioaugmentation with KB-1^®^ (TCE/ME_2001_SiREM) at two TCE-contaminated sites: one in Canada (ISSO) and another site in the UK (SaBRE). At the UK site, the abundance of *Geobacter* 16S rRNA and *rdh* genes were also monitored.

The results from the Canadian ISSO site revealed that the two most abundant *rdhA* genes in native populations (monitored prior to any treatment) were orthologs to KB1-4 and KB1-6/*bvcA,* but that their concentrations were barely above detection limits, at approximately 3x10^5^ copies per L (Table S8; Supporting Information). Although the *bvcA*-encoded dehalogenase is known to catalyze the dechlorination of cDCE all the way to ethene (63), there was little dechlorination beyond cDCE before electron donor amendment (22). During biostimulation with electron donor prior to bioaugmentation, *D. mccartyi* and *vcrA* copy numbers increased significantly, reaching a *vcrA/16S* ratio near one (Figure 4a), indicating growth of *vcrA*-bearing native populations. The abundance of *rdhA* genes orthologous to KB1-11, KB1-15, KB1-18 also increased during this period (Figure 3a). After bioaugmentation with KB-1, the *rdhA* profile did not change substantially, even though higher ethene concentrations and faster dechlorination were observed (Figure 4a). A reduction of the *bvcA/16S* ratio from ∼0.5 to 0.1 was the most notable change during this period (Figure 4a). The native populations that grew upon electron donor addition (prior to inoculation) harbored orthologs of 6 out of the 12 KB-1 *rdhA* genes monitored. The KB-1 culture originated from a site in Ontario in the Kitchener/Waterloo area within the same watershed as the ISSO site roughly100 km apart, and possibly share common microbial populations. From the data after bioaugmentation, it is clear that the dominant *D. mccartyi* populations from the KB-1 inoculum were not the ones responsible for the enhanced ethene production observed because KB1-12 and KB1-23 sequences that were dominant in the inoculum were not enriched in samples from the site. Perhaps a low-abundance *D. mccartyi* population in the inoculum grew at the site. Alternatively, bioaugmentation may have provided inoculation with supporting organisms to enhance dechlorination of VC by the native *D. mccartyi.* This result is in agreement with previous analyses of data from this site (22) that suggested that growth of a *Bacteroidetes* population present in KB-1^®^ capable of ethanol fermentation and vitamin production enhanced ethenogenesis after bioaugmentation.

**Figure 4.**
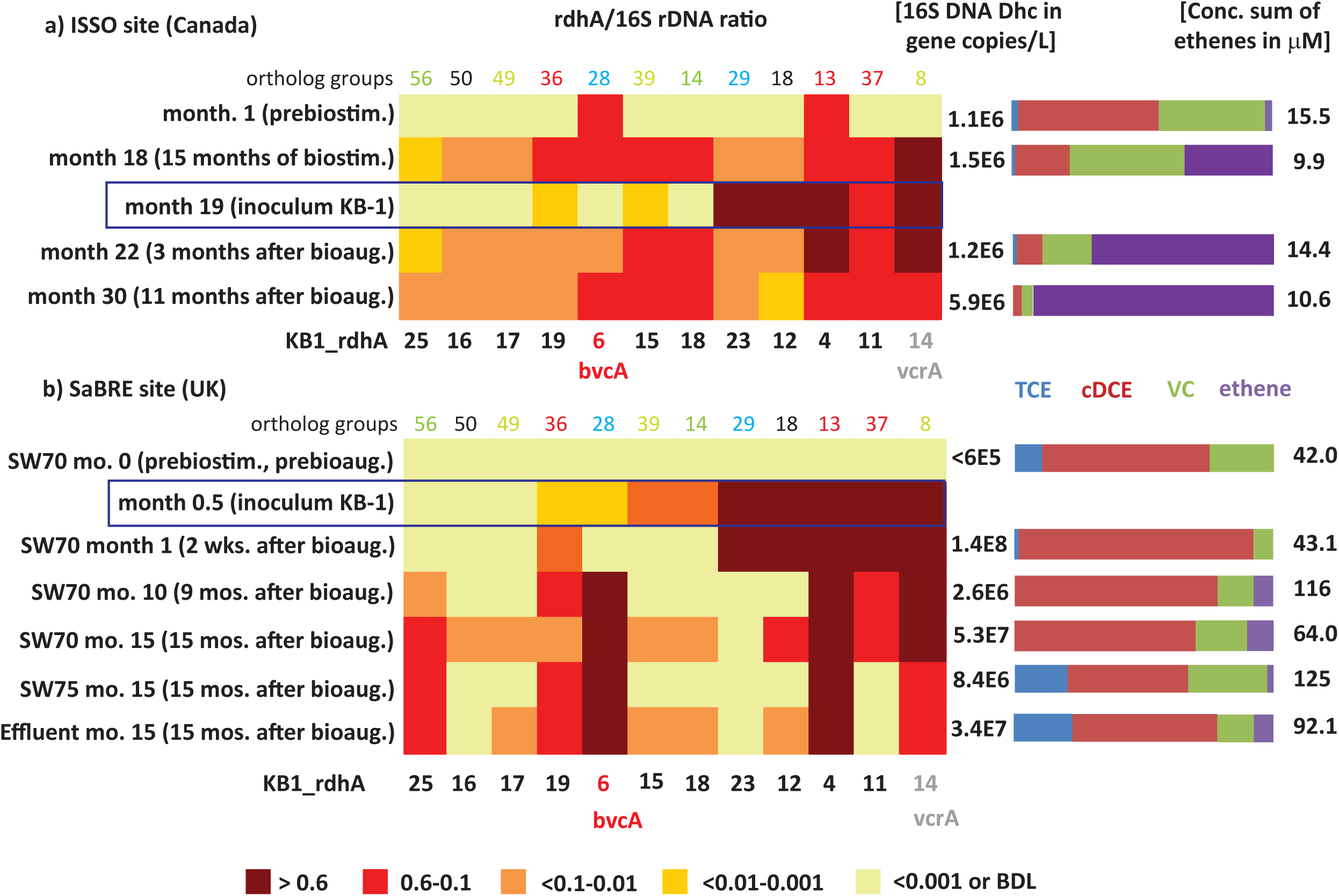
Heatmaps showing *D. mccartyi rdhA*/16S rDNA gene copy ratios in field samples. a) Canadian ISSO site. Panel b) UK SaBRE site. Corresponding absolute copies of *D. mccartyi* per L and concentrations of chlorinated ethenes are represented on the right side of each panel. Data from the ISSO site are from the composite pipeline (see Fig. S1 of Supporting Information). Data from the SaBRE site are from the influent (INF), sampling wells (SW70 and SW75) and effluent (EFF) of the test plot (See Figure S2 of Supporting Information).

The *rdhA* survey also revealed *D. mccartyi* population shifts as dechlorination progressed in the field (Figure 4a). Specifically, prior to any active treatment and when cDCE was the dominant cVOC, *bvcA* and KB1-4 were the most abundant *rdhA* genes found, a feature only seen with the cDCE enrichments in the KB-1 subculture survey. During biostimulation with electron donor, there was a gradual decrease in cDCE concentration and an increase in the concentration of VC and ethene. After 15 months of biostimulation (Month 18), the relative abundance of several other *rdhA* genes increased to above 10%, including genes similar to KB1-11, KB1-15, KB1-18, and especially *vcrA (*Figure 4a). This parttern of rdhA genes was also seen in the KB-1 VC enrichments. Three months after bioaugmentation (Month 22), the proportion of ethene relative to VC and cDCE further increased. Here, the *bvcA* ratio decreased to less than 10% while the *vcrA* ratio remained at approximately 1. At Month 30, when there was almost no VC or cDCE left, the *vcrA* ratio returned to below 0.6. At this time, relative abundance of KB1-12 decreased substantially; these *rdhA* was also found to vary between enrichment cultures.

Ratios describe shifts in the underlying *D. mccartyi* populations, but the absolute abundance of *D. mccartyi* is more relevant to observed dechlorinating activity. Major changes in absolute abundance of *rdhA* genes KB1-11, KB1-15, KB1-18 and KB1-14/*vcrA* were observed as well, starting from below detection limit (< ∼10^5^ copies L^-1^) up to 0.9-3x10^6^ copies L^-1^ (Table S8). The abundances of *vcrA* and *D. mccartyi* 16S rRNA genes increased up to the end of the monitoring period (Month 30) reaching concentrations of 1.6x10^6^ and 6x10^6^ copies L^-1^, respectively. For the *rdhA* genes KB1-12 and KB1-23, characteristic of the dominant *D. mccartyi* populations in the inoculum (Figure 4a), there was no such significant change, further indicating that this population did not contribute to the enhanced dechlorination observed after bioaugmentation.

The *rdhA* gene suite also was used to investigate groundwater from the SaBRE bioaugmentation trial within a test cell at a TCE-contaminated site in the U.K. Prior to any treatment, *D. mccartyi* gene copy numbers were below the detection limit (<6x10^4^ gene copies L^-1^) in all wells studied except the effluent (EFF) location where *D. mccartyi* titers of about 10^5^ gene copies L^-1^ were detected (data not shown). No significant VC and ethene concentrations were measured in these wells prior to treatment. Unfortunately, no samples were collected during biostimulation prior to inoculation of KB-1 (all data provided in Table S8). Two weeks after bioaugmentation with KB-1 and about a month after a single donor addition event, *D. mccartyi* titers increased by three orders of magnitude in SW70, reaching 10^8^ gene copies L^-1^ (Table S8). In a sample from this well, all KB-1 biomarkers characteristic of the original inoculum, that is KB1-4, KB1-11, KB1-12, KB1-14/*vcrA*, and KB-23 were found to have *rdhA* ratios greater than 0.6 (Figure 4b). The KB-1 *Geobacter* 16S rRNA gene and the *Geobacter rdhA* gene were also detected at significant titers in the range 0.8-4.6x10^7^ copies L^-1^ (Table S8). These data indicate that the dominant KB-1 *D. mccartyi* populations in the inoculum grew within the test cell, at least in the vicinity of this particular well (SW70).

In samples from SW-70 taken 10 and 15 months after donor addition, the ratios of KB1-11, KB1-12 and KB1-23, abundant in the inoculum and at one month, gradually decreased from values greater than 0.6 down to 0.1 (Figure 4b). The decrease of the ratio for KB1-23 was even more pronounced with a conservative estimate of 0.001 at month 15. Over the same time, the *vcrA* ratio remained greater than 0.6, while the *bvcA* ratio increased from less than 0.01 to about one (Figure 4b), indicating a dramatic shift in the dominant *D. mccartyi* populations with time in this well. A feature of this site was the consistently high concentrations of cDCE that seems to have led to the growth of *bvcA*-containing strains. At month 15, downgradient sampling locations SW75 and EFF were dominated by KB1-6/*bvcA*, KB1-4 and KB1-14/*vcrA*, and were similar to well SW70. At the influent (INF) sample location, upgradient of where the inoculum was added, gene copy numbers of *D. mccartyi* and *rdhA* genes were all below their detection limits. Curiously, the *rdhA* and 16S rRNA genes from *Geobacter* were were detected at levels just above the detection.

The Canadian ISSO and UK SaBRE sites differed in many ways. The former Ontario site had one order of magnitude lower cVOC concentrations, more complete dechlorination to ethene, a more-readily fermentable substrate (ethanol vs emulsified oil) and a recirculation system that provided better mixing of substrates and microorganisms. The SaBRE site had higher concentrations of VOCs, giving rise to higher microbial and ethene concentrations than at the Canadian site, although the dominant compound at the site was cDCE. Particularly interesting at the SaBRE site was the co-presence of both *vcrA* (KB1-14) and *bvcA* (KB1-6) after month 10 at SW70 and after month 15 at both SW75 and Effluent. Co-presence of *bvcA* and *vcrA* at field sites has been reported in various studies, with greater abundance of *bvcA* over *vcrA* at some sites and vice versa at others (24, 64, 65). Such differences have been attributed to redox potential, with *bvcA* perhaps more abundant under less reducing conditions (64). The redox potential at SaBRE was relatively low and consistent over the length of the cell and the experimental time. Based on our laboratory cultures enriched with cDCE, it seems more likely that the high concentrations of cDCE and VC at a site exerted significant selective pressure on *D. mccartyi* populations harboring different *rdhA* genes, mimicking what we also observed in enrichment cultures.

It is clear that tracking specific *rdhA* fingerprints is challenging due to growth of minor populations of *D. mccartyi* either from native or inoculum-derived strains. In future studies, it will be necessary to first identify site-specific *rdhA* genes that are not present at all in bioaugmentation cultures like KB-1, to track growth of indigenous populations. Perhaps a screening approach such as outlined in Hug *et al*. (50) based on a set of 46 primer pairs to all known *rdhA* sequences, or the approach by Mayer-Blackwell et al. (58) should be used to identify *rdh* genes in native strains prior to bioaugmentation. Then subsequent screens may be more discriminating. More significantly, this study found clear selective pressure on *D. mccartyi* populations from the most abundant terminal chlorinated electron acceptor present. In both field samples and in well-established enrichment cultures, we found that *D. mccartyi* are not a homogeneous population but rather a complex and diverse mixture of populations harbouring different complements of rdhA genes that respond and perhaps adapt selectively to external conditions - in particular electron acceptor - as revealed herein by monitoring *rdhA*/*Dhc*16S ratios. The success of biostimulation and bioaugmentation approaches relies on the growth of populations with enzymes that actively convert chlorinated ethenes past VC all the way to ethene. It seems that specific *rdhA* sequences confer populations with fitness advantages depending on their local environment, and suggests that bioaugmenting with more refined cultures acclimated to intermediates like cDCE or VC that tend to accumulate at some sites may help to overcome stall and achieve complete dechlorination at some contaminated sites. This study also underpins the need to further understand the role of horizontal gene transfer in these dechlorinating communities. These *rdhA* genes are often situated in regions of the genome that suggest potential for mobilization; yet we have no idea how quickly and under what conditions *rdhA* gene transfer occurs in organohalide-respiring bacteria such as *Dehalococcoides*.

## Supporting Information

Summary of cultures (Table S1); Summary of *rdhA* genes in KB-1 (Table S2); Timeline and sketch of the ISSO site in Canada (Figure S1); Timeline and sketch of the SaBRE site in the UK (Figure S2); Phylogenetic analysis of cultures (Text S1); Clustering analysis of *rdhA* genes (Text S2); Phylogenetic tree of *rdhA* genes based on nucleotide sequences (Figure S3); Primer pairs for qPCR quantification (Table S3); Mismatches between primers and sequences from OG groups (Table S4); qPCR standard curves (Table S5); qPCR data for Figures 2 and 3 (Tables S6-S8). Detection limits for qPCR (Table S9).

## Acknowledgements

Dr. Alfredo Pérez-de-Mora thanks the European Commission for financial support through an IOF Marie Curie Fellowship within the project AnDeMic (Contract 23974/7th Framework). The authors also wish to thank the Natural Sciences and Engineering Council of Canada (NSERC), the US Department of Defense Strategic Environmental Research and Development Program (SERDP), Genome Canada and the Ontario Genomics Institute (2009-OGI-ABC-1405), the Government of Ontario through the ORF-GL2 program and Sustainable Development Technology Canada (SDTC) for providing funding. The authors also acknowledge R&D in kind contributions by Geosyntec and SiREM to the project study. The authors wish to thank Adria Bells, Jeff Roberts and Carey Austrins for field assistance and Olivia Molenda for a thorough review of the manuscript.

